# Improving gene-network inference with graph-wavelets and making insights about ageing associated regulatory changes in lungs

**DOI:** 10.1101/2020.07.24.219196

**Authors:** Shreya Mishra, Divyanshu Srivastava, Vibhor Kumar

## Abstract

Using gene-regulatory-networks based approach for single-cell expression profiles can reveal unprecedented details about the effects of external and internal factors. However, noise and batch effect in sparse single-cell expression profiles can hamper correct estimation of dependencies among genes and regulatory changes. Here we devise a conceptually different method using graph-wavelet filters for improving gene-network (GWNet) based analysis of the transcriptome. Our approach improved the performance of several gene-network inference methods. Most Importantly, GWNet improved consistency in the prediction of generegulatory-network using single-cell transcriptome even in presence of batch effect. Consistency of predicted gene-network enabled reliable estimates of changes in the influence of genes not highlighted by differential-expression analysis. Applying GWNet on the single-cell transcriptome profile of lung cells, revealed biologically-relevant changes in the influence of pathways and master-regulators due to ageing. Surprisingly, the regulatory influence of ageing on pneumocytes type II cells showed noticeable similarity with patterns due to effect of novel coronavirus infection in Human Lung.

## Introduction

Inferring gene-regulatory-networks and using them for system-level modelling is being widely used for understanding the regulatory mechanism involved in disease and development. The interdependencies among variables in the network is often represented as weighted edges between pairs of nodes, where edge weights could represent regulatory interactions among genes. Gene-networks can be used for inferring causal models [1], designing and understanding perturbation experiments, comparative analysis [2] and drug discovery [3]. Due to wide applicability of network inference, many methods have been proposed to estimate interdependencies among nodes. Most of the methods are based on pairwise correlation, mutual information or other similarity metrics among gene expression values, provided in a different condition or time point. However, resulting edges are often influenced by indirect dependencies owing to low but effective background similarity in patterns. In many cases, even if there is some true interaction among a pair of nodes, its effect and strength is not estimated properly due to noise, background-pattern similarity and other indirect dependencies. Hence recent methods have started using alternative approaches to infer more confident interactions. Such alternative approach could be based on partial correlations [4] or ARACNE’s method of statistical threshold of mutual information [5].

Single-cell expression profiles often show heterogeneity in expression values even in a homogeneous cell population. Such heterogeneity can be exploited to infer regulatory networks among genes and identify dominant pathways in a celltype. However, due to the sparsity and ambiguity about the distribution of gene expression from single-cell RNA-seq profiles, the optimal measures of gene–gene interaction remain unclear. Hence recently, Sknnider et al. [6] evaluated 17 measures of association to infer gene co-expression based network. In their analysis, they found two measures of association, namely phi and rho as having the best performance in predicting co-expression based gene-gene interaction using scRNA-seq profiles. In another study, Chen et al. [7] performed independent evaluation of a few methods proposed for genenetwork inference using scRNA-seq profiles such as SCENIC [8], SCODE [9], PIDC [10]. Chen et al. found that for single-cell transcriptome profiles either generated from experiments or simulations, these methods had a poor performance in reconstructing the network. Performance of such methods can be improved if gene-expression profiles are denoised. Thus the major challenge of handling noise and dropout in scRNA-seq profile is an open problem. The noise in single-cell expression profiles could be due to biological and technical reasons. The biological source of noise could include thermal fluctuations and a few stochastic processes involved in transcription and translation such as allele specific expression [11] and irregular binding of transcription factors to DNA. Whereas technical noise could be due to amplification bias and stochastic detection due to low amount of RNA. Raser and O’Shea [12] used the term noise in gene expression as measured level of its variation among cells supposed to be identical. Raser and O’Shea categorised potential sources of variation in gene-expression in four types: (i) the inherent stochasticity of biochemical processes due to small numbers of molecules; (ii) heterogeneity among cells due to cell-cycle progression or a random process such as partitioning of mitochondria (iii) subtle micro-environmental differences within a tissue (iv) genetic mutation. Overall noise in gene-expression profiles hinders in achieving reliable inference about regulation of gene activity in a cell-type. Thus, there is demand for pre-processing methods which can handle noise and sparsity in scRNA-seq profiles such that inference of regulation can be reliable.

The predicted gene-network can be analyzed further to infer salient regulatory mechanisms in a cell-type using methods borrowed from Graph theory. Calculating gene-importance in term of centrality, finding communities and modules of genes are common downstream analysis procedures [2]. Just like gene-expression profile, inferred gene network could also be used to find differences in two groups of cells(sample) [13] to reveal changes in the regulatory pattern caused due to disease, environmental exposure or ageing. In particular, a comparison of regulatory changes due to ageing has gained attention recently due to a high incidence of metabolic disorder and infection based mortality in the older population. Especially in the current situation of pandemics due to novel coronavirus (SARS-COV-2), when older individuals have a higher risk of mortality, a question is haunting researchers. That question is: Why old lung cells have a higher risk of developing severity due to SARS-COV-2 infection. However, understanding regulatory changes due to ageing using gene-network inference with noisy single-cell scRNA-seq profiles of lung cells is not trivial. Thus there is a need of a noise and batch effect suppression method for investigation of the scRNA-seq profile of ageing lung cells [14] using a network biology approach.

Here we have developed a method to handle noise in gene-expression profiles for improving gene-network inference. Our method is based on graph-wavelet based filtering of gene-expression. Our approach is not meant to overlap or compete with existing network inference methods but its purpose is to improve their performance. Hence, we compared other output of network inference methods with and without graph-wavelet based pre-processing. We have evaluated our approach using several bulk sample and single-cell expression profiles. We further investigated how our denoising approach influences the estimation of graph-theoretic properties of gene-network. We also asked a crucial question: how the gene regulatory-network differs between young and old individual lung cells. Further, we compared the pattern in changes in the influence of genes due to ageing with differential expression in COVID infected lung.

## Results

Our method uses a logic that cells (samples) which are similar to each other, would have a more similar expression profile for a gene. Hence, we first make a network such that two cells are connected by an edge if one of them is among the top K nearest neighbours (KNN) of the other. After building KNN-based network among cells (samples), we use graph-wavelet based approach to filter expression of one gene at a time (see Fig. 1). For a gene, we use its expression as a signal on the nodes of the graph of cells. We apply a graph-wavelet transform to perform spectral decomposition of graph-signal. After graph-wavelet transformation, we choose the threshold for wavelet coefficients using sureShrink and BayesShrink or a default percentile value determined after thorough testing on multiple data-sets. We use the retained values of the coefficient for inverse graph-wavelet transformation to reconstruct a filtered expression matrix of the gene. The filtered gene-expression is used for gene-network inference and other down-stream process of analysis of regulatory differences. For evaluation purpose, we have calculated inter-dependencies among genes using 5 different co-expression measurements, namely Pearson and spearman correlations, *ϕ* and *ρ* scores and ARACNE.

**Figure 1:**
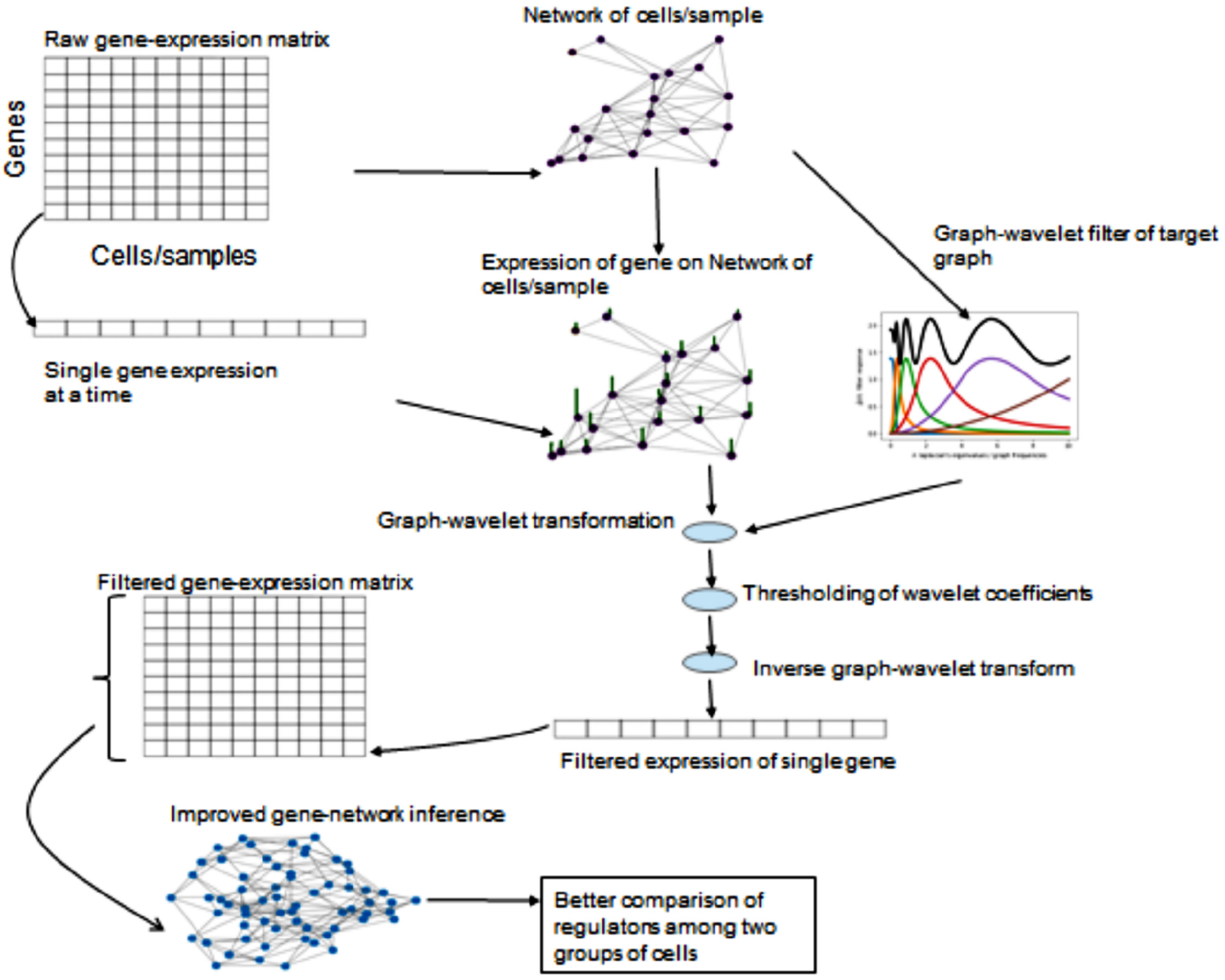
The flowchart of GWNet pipeline. First, a KNN based network is made between samples/cell. A filter for graph wavelet is learned for the KNN based network of samples/cells. Gene-expression of one gene at a time is filtered using graph-wavelet transform. Filtered gene-expression data is used for network inference. The inferred network is used to calculate centrality and differential centrality among groups of cells.

### Evaluation using bulk expression profiles from DREAM challenge

The biological and technical noise can both exist in a bulk sample expression profile ([12]). In order to test the hypothesis that graph-based denoising could improve gene-network inference, we first evaluated the performance of our method on bulk expression data-set. We used 4 data-sets made available by DREAM5 challenge consortium [15]. Three data-sets were based on the original expression profile of bacterium Escherichia coli and the single-celled eukaryotes Saccharomyces cerevisiae and S aureus. While the fourth data-set was simulated using in silico network with the help of GeneNetWeaver, which models molecular noise in transcription and translation using chemical Langevin equation [16]. The true positive interactions for all the four data-sets are also available. We compared Graph Fourier based low pass-filtering with graph-wavelet based denoising using three different approaches to threshold the wavelet-coefficients. We achieved 5 - 25 % improvement in score over raw data based on DREAM5 criteria [15] with correlation, ARACNE and rho based network prediction. With *ϕ_s_* based gene-network prediction, there was an improvement in 3 out of 4 DREAM5 data-sets (Fig. 2A). All the 5 network inference methods showed improvement after graph-wavelet based denoising of simulated data (in silico) from DREAM5 consortium (Fig. 2A). Moreover, graph-wavelet based filtering had better performance than Chebyshev filter-based low pass filtering in graph Fourier domain. It highlights the fact that even bulk sample data of gene-expression can have noise and denoising it with graph-wavelet after making KNN based graph among samples has the potential to improve gene-network inference. Moreover, it also highlights another fact, well known in the signal processing field, that wavelet-based filtering is more adaptive than low pass-filtering.

**Figure 2:**
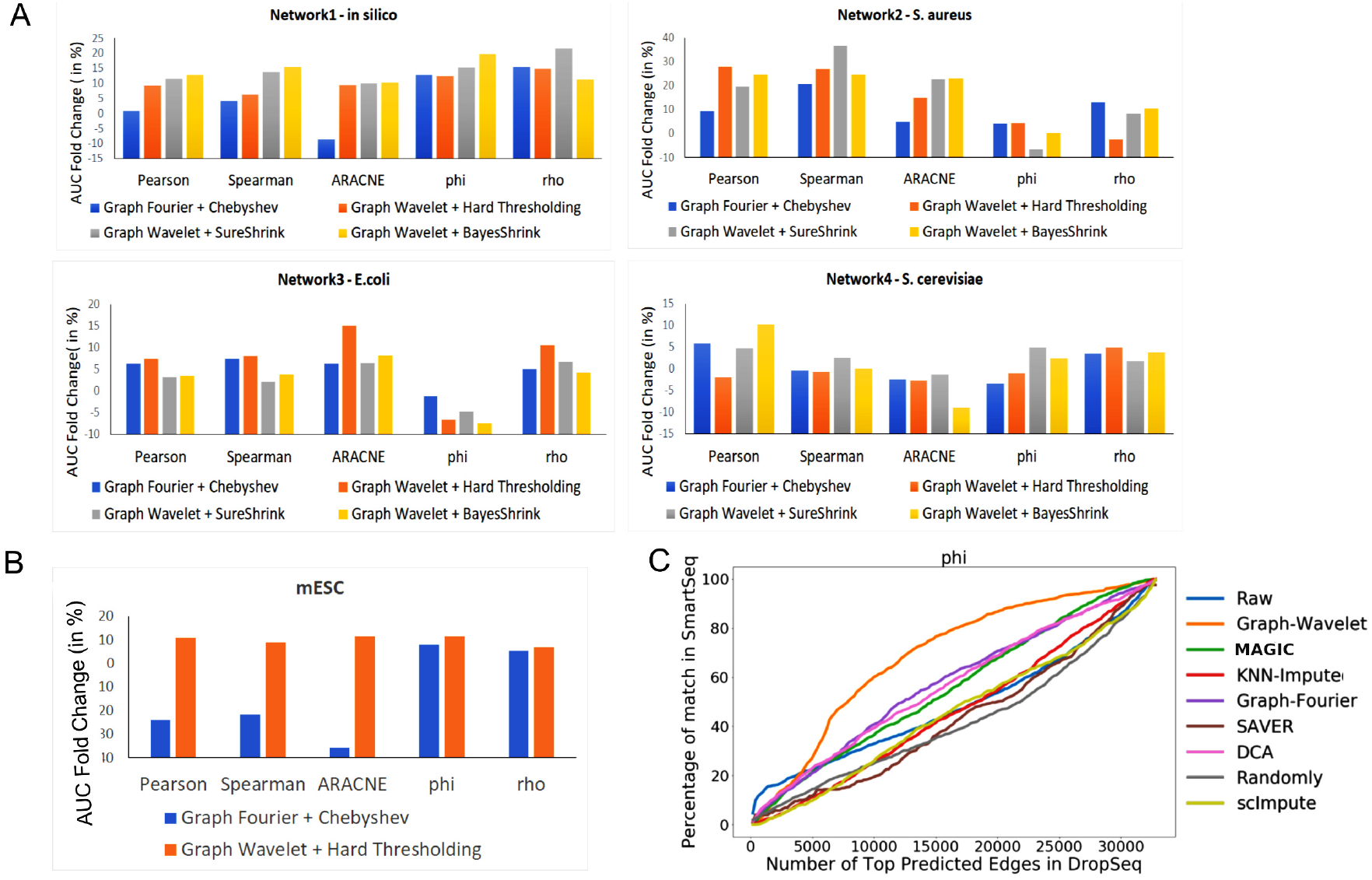
Improvement in gene-network inference by graph-wavelet based denoising of gene-expression (A) Performance of network inference methods using bulk gene-expression data-sets of DREAM5 challenge. Three different ways of shrinkage of graph-wavelet coefficients were compared to graph-Fourier based low pass filtering. The Y-axis shows fold change in area under curve(AUC) for receiver operating characteristic curve (ROC) for overlap of predicted network with golden-set of interactions. For hard threshold, the default value of 70% percentile was used. (B) Performance evaluation using single-cell RNA-seq (scRNA-seq) of mouse embryonic stem cells (mESCs) based network inference after filtering the gene-expression. The gold-set of interactions was adapted from [19] (C) Comparison of graph wavelet-based denoising with other related smoothing and imputing methods in terms of consistency in the prediction of the gene-interaction network. Here, Phi (*ϕ_s_*) score was used to predict network among genes. For results based on other types of scores see supplementary Figure S1. Predicted networks from two scRNA-seq profile of mESC were compared to check robustness towards the batch effect.

### Graph-wavelet based denoising of single-cell expression profiles improves gene-networks inference

In comparison to bulk samples, there is a higher level of noise and dropout in single-cell expression profiles. Dropouts are caused by non-detection of true expression due to technical issues. Using low-pass filtering after graph-Fourier transform seems to be an obvious choice as it fills in a background signal at missing values and suppresses high-frequency outlier-signal [17]. However, in the absence of information about cell-type and cellstates, a blind smoothing of a signal may not prove to be fruitful. Hence we applied graph-wavelet based filtering for processing gene-expression data-set from the scRNA-seq profile. We first used scRNA-seq data-set of mouse Embryonic stem cells (mESCs) [18]. In order to evaluate network inference in an unbiased manner, we used gene regulatory interactions compiled by another research group [19]. Our approach of graph-wavelet based pre-processing of mESC scRNA-seq data-set improved the performance of gene-network inference methods by 8-10 percentage (Fig. 2B). However, most often, the gold-set of interaction used for evaluation of gene-network inference is incomplete, which hinders the true assessment of improvement.

Hence we also used another approach to validate our method. For this purpose, we used a measure of overlap among network inferred from two scRNA-seq data-sets of the same cell-type but having different technical biases and batch effects. If the inferred networks from both data-sets are closer to true gene-interaction model, they will show high overlap. For this purpose, we used two scRNA-seq data-set of mESC generated using two different protocols(SMARTseq and Drop-seq). For comparison of consistency and performance, we also used a few other imputation and denoising methods proposed to filter and predict the missing expression values in scRNA-seq profiles. We evaluated 7 other such methods; Graph-Fourier based filtering [17], MAGIC [20], scImpute [21], DCA [22], SAVER [23], Randomly [24], KNN-impute [25]. Graph-wavelet based denoising provided better improvement in AUC for overlap of predicted network with known interaction than other 7 methods meant for imputing and filtering scRNA-seq profiles (supplementary Figure S1A). Similarly in comparison to graph-wavelet based denoising, the other 7 methods did not provided substantial improvement in AUC for overlap among gene-network inferred by two data-sets of mESC (Fig. 2C, supplementary Figure S1B). However, graph wavelet-based filtering improved the overlap between networks inferred from different batches of scRNA-seq profile of mESC even if they were denoised separately (Fig. 2C, supplementary Figure S1B). With *ϕ_s_* based edge scores the overlap among predicted gene-network increased by 80% due to graph-wavelet based denoising (Fig. 2C). The improvement in overlap among networks inferred from two batches hints that graph-wavelet denoising is different from imputation methods and has the potential to substantially improve gene-network inference using their expression profiles.

### Improved gene-network inference from single-cell profile reveal age-based regulatory differences

Improvement in overlap among inferred gene-networks from two expression data-set for a cell type also hints that after denoising predicted networks are closer to true gene-interaction profiles. Hence using our denoising approach before estimating the difference in inferred gene-networks due to age or external stimuli could reflect true changes in the regulatory pattern. Such a notion inspired us to compare gene-networks inferred for young and old pancreatic cells using their scRNA-seq profile filtered by our tool [26]. Martin et al. defined three age groups, namely juvenile (1month-6 years), young adult (21-22 years) and aged (38-54 years) [26]. We applied graph-wavelet based denoising of pancreatic cells from three different groups separately. In other words, we did not mix cells from different age groups while denoising. Graph-wavelet based denoising of a single-cell profile of pancreatic cells caused better performance in terms of overlap with protein-protein interaction (PPI) (Fig. 3A, Supplementary Figure S2A). Even though like Chen et al. [7] we have used PPI to measure improvement in gene-network inference, it may not be reflective of all gene-interactions. Hence we also used the criteria of increase in overlap among predicted networks for same cell-types to evaluate our method for scRNA-seq profiles of pancreatic cells. Denoising scRNA-seq profiles also increased overlap between inferred gene-network among pancreatic cells of the old and young individuals (Fig. 3B, Supplementary Figure S2B). We performed quantile normalization of original and denoised expression matrix taking all 3 age groups together to bring them on the same scale to calculate the variance of expression across cells of every gene. The old and young pancreatic alpha cells had a higher level of median variance of expression of genes than juvenile. However, after graph-wavelet based denoising, the variance level of genes across all the 3 age groups became almost equal and had similar median value (Fig. 3C). Notice that, it is not trivial to estimate the fraction of variances due to transcriptional or technical noise. Nonetheless, graph-wavelet based denoising seemed to have reduced the noise level in single-cell expression profiles of old and young adults. Differential centrality in the co-expression network has been used to study changes in the influence of genes. However, noise in single-cell expression profiles can cause spurious differences in centrality. Hence we visualized the differential degree of genes in network inferred using young and old cells scRNA-seq profiles. The networks inferred from non-filtered expression had a much higher number of non-zero differential degree values in comparison to the de-noised version (Fig. 3D, Supplementary Figure S2C). Thus denoising seems to reduce differences among centrality, which could be due to randomness of noise. Next, we analyzed the properties of genes whose variance dropped most due to graph-wavelet based denoising. Surprisingly, we found that top 500 genes with the highest drop in variance due to denoising in old pancreatic beta cells were significantly associated with diabetes Mellitus and hyperinsulinism. Whereas, top 500 genes with the highest drop in variance in young pancreatic beta cells had no or insignificant association with diabetes (Fig. 3E). A similar trend was observed with pancreatic alpha cells (Supplementary Figure S2D). Such a result hint that ageing causes increase in stochasticity of the expression level of genes associated with pancreas function and denoising could help in properly elucidating their dependencies with other genes.

**Figure 3:**
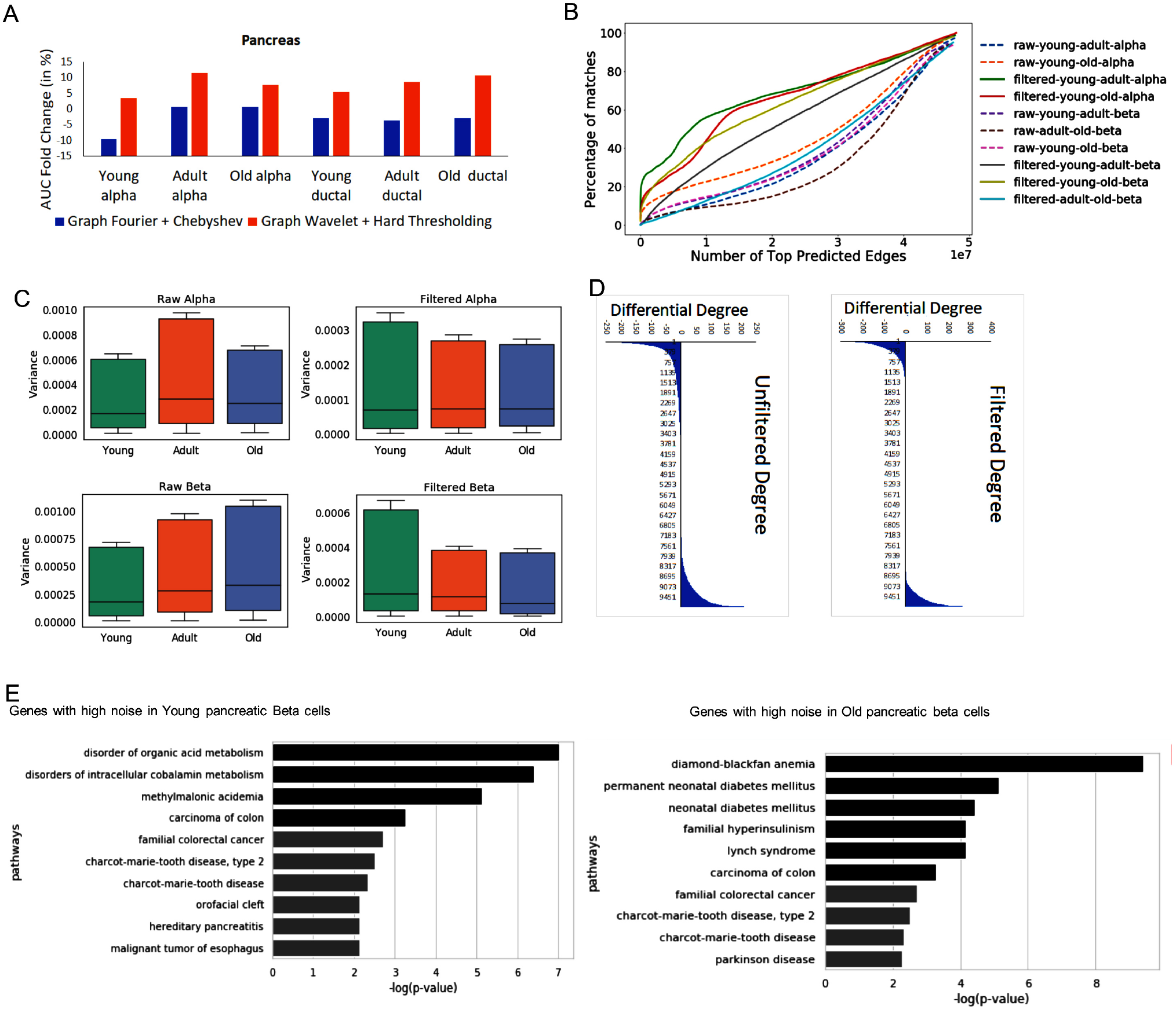
Performance and analysis of noise for single-cell RNA-seq profile of pancreatic cells. (A) Performance based on overlap of predicted network with protein-protein interaction data-set. (B) Evaluation of consistency of predicted network. For comparing two networks it is important to reduce differences due to noise. Hence the plot here shows similarity of predicted networks before and after graph-wavelet based denoising. The result shown here are for correlation-based co-expression network, while similar results are shown using *ρ* score in supplementary Figure S2. (C) Variances of expression of genes across single-cells before and after denoising (filtering) is shown here. Variances of genes in a cell-type was calculated separately for 3 different stages of ageing (young, adult and old). The variance (estimate of noise) is higher in older alpha and beta cells compared to young. However, after denoising variance of genes in all ageing stage becomes equal (D) Effect of noise in estimated differential centrality is shown is here. The difference in the degree of genes in network estimated for old and young pancreatic beta cells is shown here. The number of non-zero differential-degree estimated using denoised expression is lower than unfiltered expression based networks.(E) Enriched panther pathway terms for top 500 genes with the highest drop in variance after denoising in old and young pancreatic beta cells.

### Improvement in Gene-network inference for studying regulatory differences among young and old lung cells

Studying cell-type-specific changes in regulatory networks due to ageing has the potential to provide better insight about predisposition for disease in the older population. Hence we inferred gene-network for different cell-types using scRNA-seq profiles of young and old mouse lung cells published by Kimmel et al. [14].The lower lung epithelia where a few viruses seem to have the most deteriorating effect consists of multiple types of cells such as bronchial epithelial and alveolar epithelial cells, fibroblast, alveolar macrophages, endothelial and other immune cells. The alveolar epithelial cells, also called as pneumocytes are of two major types. The type 1 alveolar (AT1) epithelial cells for major gas exchange surface of lung alveolus has an important role in the permeability barrier function of the alveolar membrane. Type 2 alveolar cells (AT2) are the progenitors of type 1 cells and has the crucial role of surfactant production. AT2 cells (or pneumocytes type II) cells are a prime target of many viruses; hence it is important to understand the regulatory patterns in AT2 cells, especially in the context of ageing.

We applied our method of denoising on scRNA-seq profiles of cells derived from old and young mice lung [14]. Graph wavelet based denoising lead to an increase in consistency among inferred gene-network for young and old mice lung for multiple cell-types (Fig. 4A). Graph-wavelet based denoising also lead to an increase in consistency in predicted gene-network from data-sets published by two different groups (Fig. 4B). The increase in overlap of gene-networks predicted for old and young cells scRNA-seq profile, despite being denoised separately, hints about a higher likelihood of predicting true interactions. Hence the chances of finding gene-network based differences among old and young cells were less likely to be dominated by noise. We studied ageing-related changes in PageRank centrality of nodes(genes). Since PageRank centrality provides a measure of “popularity” of nodes, studying its change has the potential to highlight the change in the influence of genes. First, we calculated differential PageRank of genes among young and old AT2 cells (supporting File-1) and performed gene-set enrichment analysis using Enrichr [27]. The top 500 genes with higher PageRank in young AT2 cells had enriched terms related to integrin signalling, 5HT2 type receptor mediated signalling, H1 histamine receptor-mediated signalling pathway, VEGF, cytoskeleton regulation by Rho GTPase and thyrotropin activating receptor signalling (Fig. 4C). We ignored oxytocin and thyrotropin-activating hormone-receptor mediated signalling pathways as an artefact as the expression of oxytocin and TRH receptors in AT2 cells was low. Moreover, genes appearing for the terms “oxytocin receptor-mediated signalling” and “thyrotropin activating hormone-mediated signalling” were also present in gene-set for 5Ht2 type receptormediated signalling pathway. We found literature support for activity in AT2 cells for most of the enriched pathways. However, there were very few studies which showed their differential importance in old and young cells, such as Bayer et al. demonstrated mRNA expression of several 5-HTR including 5-Ht2, 5Ht3 and 5Ht4 in alveolar epithelial cells type II (AT2) cells and their role in calcium ion mobilization. Similarly, Chen et al. [28] showed that histamine 1 receptor antagonist reduced pulmonary surfactant secretion from adult rat alveolar AT2 cells in primary culture. VEGF pathway is active in AT2 cells, and it is known that ageing has an effect on VEGF mediated angiogenesis in lung. Moreover, VEGF based angiogenesis is known to decline with age [29]. We further performed gene-set enrichment analysis for genes with increased pageRank in older mice AT2 cells. For top 500 genes with higher pageRank in old AT2 cells, the terms which appeared among 10 most enriched in both Kimmel et al. and Angelids et al. data-sets were T cell activation, B cell activation, cholesterol biosynthesis and FGF signaling pathway, angiogenesis and cytoskeletal regulation by Rho GTPase (Fig. 4D). Thus, there was 60% overlap in results from Kimmel et al. and Angelids et al. data-sets in terms of enrichment of pathway terms for genes with higher pageRank in older AT2 cells (supplementary Figure S3A, supporting file-2, supporting file-3). Overall in our analysis, inflammatory response genes showed higher importance in older AT2 cells. The increase in the importance of cholesterol biosynthesis genes hand in hand with higher inflammatory response points towards the influence of ageing on the quality of pulmonary surfactants released by AT2. Al Saedy et al. recently showed that high level of cholesterol amplifies defects in surface activity caused by oxidation of pulmonary surfactant [30].

**Figure 4:**
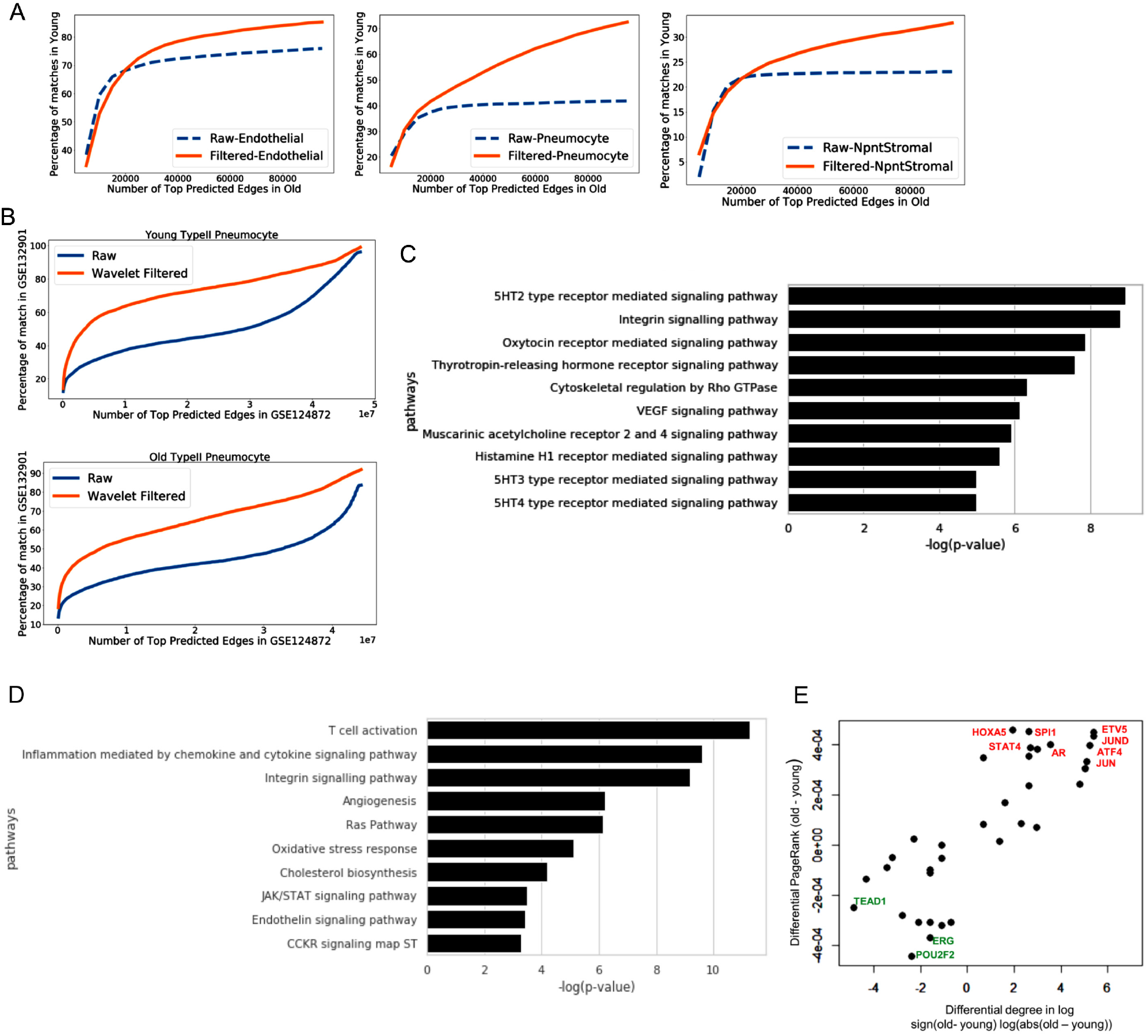
Improved regulatory inferences with graph-wavelet based pre-processing of the single-cell transcriptome of ageing lung cells (A) Consistency of prediction of the networks using the scRNA-seq profile (Kimmel et al. data-set) of young and old lung cells. The coverage of top 10000 edges in young cells in network inferred for old cells is shown here. For the same type of cells, the predicted networks for old and young cells seem to have higher overlap after graph-wavelet based filtering. The label “Raw” here means that, both networks (for old and young) were inferred using unfiltered scRNA-seq profiles. Wheres the same result from denoised scRNA-seq profile is shown as filtered. Networks were inferred using correlation-based co-expression. (B) Plot showing the overlap of networks predicted from two different data-sets with their own batch effect. X-axis shows the number of predicted edges in network predicted using Angelidis et al. data-set (GEO Id: GSE124872). Y-axis shows the fraction of top 10000 edges in network estimated using Kimmel et al. data-set. (C) 10 most enriched Panther pathway terms for top 500 genes with higher PageRank in young AT2 cells compared to old. (D) 10 most enriched panther pathway terms for top 1000 genes with higher PageRank for old AT2 cells10compared to young. (E) Scatter plot of differential degree and PageRank (old-young) of transcription factors (TF) estimated using networks predicted for old and young AT2 cells from Kimmel et al. data-set. Only TFs with a non-zero differential degree are shown.

We also performed Enrichr based analysis of differentially expressed genes in old AT2 cells (supporting File-4). For genes up-regulated in old AT2 cells compared to young, terms which reappeared were cholesterol biosynthesis, T cell and B cell activation pathways, Angiogenesis and Inflammation mediated by chemokine and cytokine signalling. Whereas few terms like RAS pathway, JAK/STAT signalling and cytoskeletal signalling by Rho GTPase did not appear as enriched for genes upregulated in old AT2 cells (Figure 3B, supporting File-4). However previously, it has been shown that the increase in age changes the balance of pulmonary renin-angiotensin system (RAS), which is correlated with aggravated inflammation and more lung injury [31]. JAK/STAT pathway is known to be involved in the oxidative-stress induced decrease in the expression of surfactant protein genes in AT2 cells [32]. Overall, these results indicate that even though the expression of genes involved in relevant pathways may not show significant differences due to ageing, but their regulatory influence could be changing substantially.

In order to further gain insight, we analyzed the changes in the importance of transcription factors in ageing AT2 cells. Among top 500 genes with higher PageRank in old AT2 cells, we found several relevant TFs. However, to make a stringent list, we considered only those TFs which had nonzero value for change in degree among gene-network for old and young AT2 cells. Overall, with Kimmel at el. data-set, we found 46 TFs with a change in PageRank and degree (supplementary table-1) due to ageing for AT2 cells (Fig. 4E). The changes in centrality (PageRank and degree) of TFs with ageing was coherent with pathway enrichment results. Such as ETV5 which has higher degree and PageRank in older cells, is known to be stabilized by RAS signalling in AT2 cells [33]. In the absence of Etv5 AT2 cell differentiate to AT1 cells [33]. Another TF Jun (c-jun) having stronger influence in old AT2 cells, is known to regulate inflammation lung alveolar cells [34]. We also found Jun to be having co-expression with Jund and Etv5 in old AT2 cell (Supplementary Figure S4). Jund whose influence seems to increase in aged AT2 cells is known to be involved in cytokine-mediated inflammation. Among the TFs Stat 1-4 which are involved in JAK/STAT signalling, Stat4 showed higher degree and PageRank in old AT2. Androgen receptor(Ar) also seem to have a higher influence in older AT2 cells (Fig. 4E). Androgen receptor has been shown to be expressed in AT2 cells [35].

We further performed a similar analysis for the scRNA-seq profile of interstitial macrophages(IMs) in lungs and found literature support for the activity of enriched pathways (supporting File-5). Whereas gene-set enrichment output for important genes in older IMs had some similarity with results from AT2 cells as both seem to have higher pro-inflammatory response pathway such as T cell activation and JAK/STAT signalling. However, unlike AT2 cells, ageing in IMs seem to cause an increase in glycolysis and pentose phosphate pathway. Higher glycolysis and pentose phosphate pathway activity levels have been previously reported to be involved in the pro-inflammatory response in macrophages by Viola et al. [36]. In our results, RAS pathway was not enriched significantly for genes with a higher importance in older macrophages. Such results show that the pro-inflammatory pathways activated due to aging could vary among different cell-types in lung.

### Comparison of the effect of ageing with COVID-19 infection in lung

In current pandemic due to SARS-COV-2, a trend has emerged that older individuals have a higher risk of developing severity and lung fibrosis than the younger population. Since our analysis revealed changes in the influence of genes in lung cells due to ageing, we compared our results with expression profiles of lung infected with SARS-COV-2 published by Blanco-Melo et al. [37]. Recently it has been shown that AT2 cells predominantly express ACE2, the host cell surface receptor for SARS-CoV-2 attachment and infection[38]. Thus COVID infection could have most of the dominant effect on AT2 cells. We found that genes with significant upregulation in SARS-COV-2 infected lung also had higher PageRank in gene-network inferred for older AT2 cells (Fig. 5a). We also repeated the process of network inference and calculating differential centrality among old and young using all types of cells in the lung together (supporting File-6). We performed gene-set enrichment for genes up-regulated in SARS-COV-2 infected lung. Majority of the 7 panther pathway terms enriched for genes up-regulated in SARS-COV-2 infected lung also had enrichment for genes with higher PageRank in old lung cells (combined). Total 6 out of 7 significantly enriched panther pathways for genes up-regulated in COVID-19 infected lung, were also enriched for genes with higher PageRank in older AT2 cells in either of the two data-sets used here (5 in Angelids et al., 3 in Kimmel et al. data-based results).

**Figure 5:**
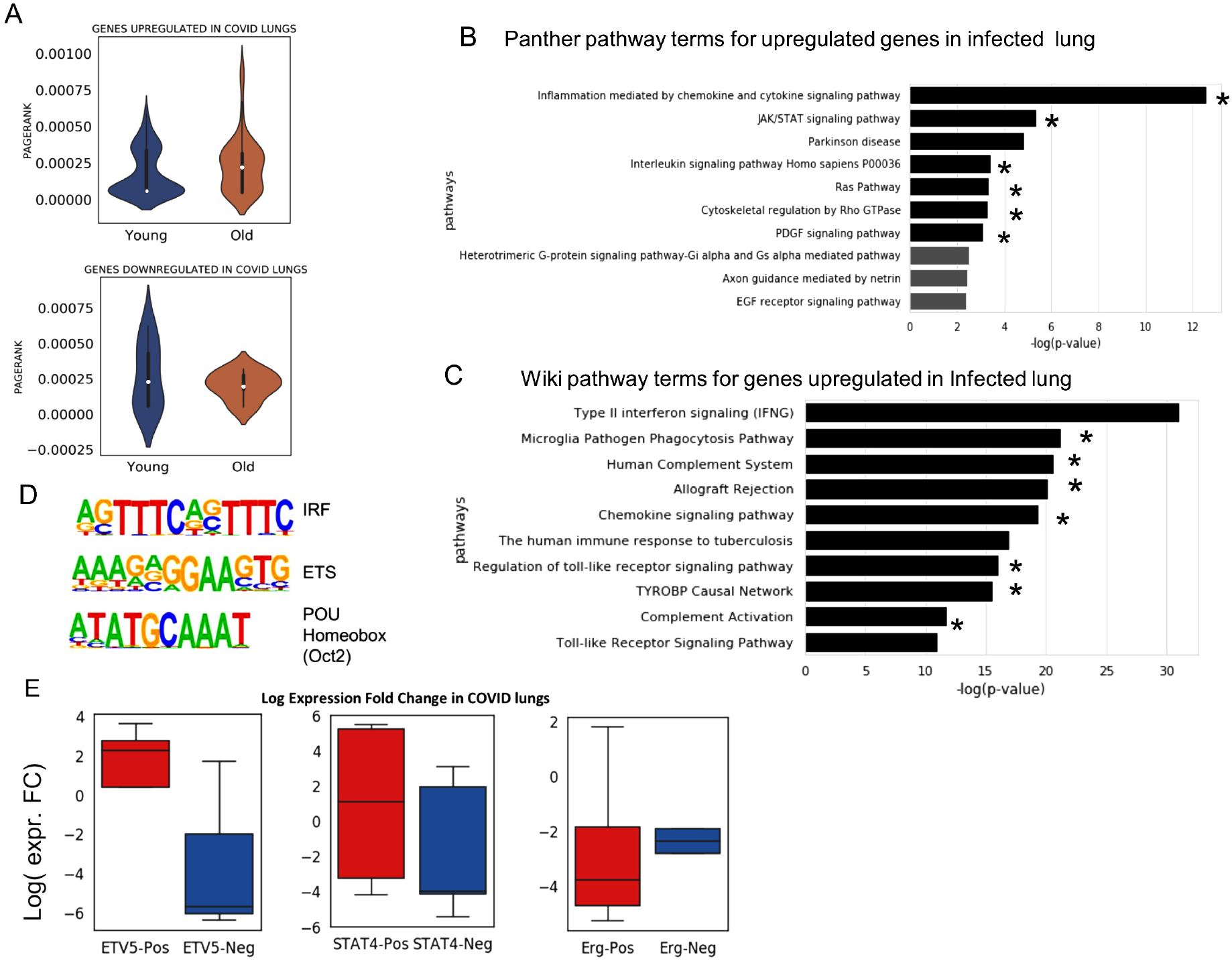
Analysis of gene-expression profile in lungs infected with SARS-COV-2 (COVID). (A) The distribution of PageRank of genes up-regulated in COVID infected lung (*FDR* < 0.05) [37]. The PageRank is shown for network estimated using the scRNA-seq profile of young and old AT2 cells. On similar pattern PageRank is shown for genes down-regulated (*FDR* < 0.05) in COVID infected lung. (B) Top 10 panther pathway enriched for genes up-regulated in COVID infected lung. The terms with * sign also have significant enrichment for genes with higher pagerRank in old AT2 cells (C) Top 10 wiki pathway terms enriched for genes up-regulated in COVID infected lung. The terms with * are also enriched (*P value* < 0.05) for genes with higher pageRank in old AT2 cells. (D) Top 3 motifs of known transcription factors (TF) enriched in promoters of genes up-regulated in COVID infected lung. (E) Fold change of expression in the lung with COVID infection for genes positively and negatively correlated with transcription factors in old AT2 cells. The results are shown for 2 transcription factors (TFs) Etv5 and Stat4, which has higher PageRank in old AT2 cells. As a control, the results are also shown for Erg, which have higher PageRank in young AT2 cells. Most of the genes which had a positive correlation with Etv5 and Stat4 expression in old murine AT2 cells were up-regulated in COVID infected lung. Whereas for Erg the trend is the opposite. Genes positively correlated with ERG genes in old AT2 had more down-regulation than genes with negative correlation. Such results hint that TFs whose influence (PageRank) increase during ageing could be involved activating or poising the genes up-regulated in COVID infection.

Among the top 10 enriched wikipathway terms for genes up-regulated in COVID infected lung, 7 has significant enrichment for genes with higher pageRank in old AT2 cells (supporting File-7). However, the term type-II interferon signalling did not have significant enrichment for genes with higher PageRank in old AT2 cells. We further investigated enriched motifs of transcription factors in promoters of genes up-regulated in COVID infected lungs (supplementary methods). For promoters of genes up-regulated in COVID infected lung top two enriched motifs belonged to IRF (interferon regulatory factor) and ETS family TFs. Notice that Etv5 belong to sub-family of ETS groups of TFs. Further analysis also revealed that most of the genes whose expression is positively correlated with Etv5 in old AT2 cells is up-regulated in COVID infected lung. In contrast, genes with negative correlation with Etv5 in old AT2 cells were mostly down-regulated in COVID infected lung. A similar trend was found for Stat4 gene. However, for Erg gene with higher pageRank in young AT2 cell, the trend was the opposite. In comparison to genes with negative correlation, positively correlated genes with Erg in old AT2 cell, had more downregulation in COVID infected lung. Such trend shows that a few TFs like Etv5, Stat4 with higher PageRank in old AT2 cells could be having a role in poising or activation of genes which gain higher expression level on COVID infection.

## Discussion

Inferring regulatory changes in pure primary cells due to ageing and other conditions, using single-cell expression profiles has tremendous potential for various applications. Such applications could be understanding the cause of development of a disorder or revealing signalling pathways and master regulators as potential drug targets. Hence to support such studies, we developed GWNet to assist biologists in work-flow for graph-theory based analysis of single-cell transcriptome. GWNet improves inference of regulatory interaction among genes using graph-wavelet based approach to reduce noise due to technical issues or cellular biochemical stochasticity in gene-expression profiles. We demonstrated the improvement in gene-network inference using our filtering approach with 4 benchmark data-sets from DREAM5 consortium and several single-cell expression profiles. Using 5 different ways for inferring network, we showed how our approach for filtering gene-expression can help gene-network inference methods. Our results of comparison with other imputation, smoothing methods and graph-Fourier based filtering showed that graph-wavelet is more adaptive to changes in the expression level of genes with changing neighborhood of cells. Thus graph-wavelet based denoising is a conceptually different approach for preprocessing of gene-expression profiles. There is a huge body of literature on inferring gene-networks from bulk gene-expression profile and utilizing it to find differences among two groups of samples. However, applying classical procedures on single-cell transcriptome profiles has not proved to be effective. Our method seems to resolve this issue by increasing consistency and overlap among gene-networks inferred using an expression from different sources (batches) for the same cell-type even if each data-sets was filtered independently. Such an increase in overlap among predicted network from independently processed data-sets from different sources hint that estimated dependencies among genes reach closer to true values after graph-wavelet based denoising of expression profiles. Having network prediction closer to true values increases the reliability of comparison of a regulatory pattern among two groups of cells. Moreover, recently Chow and Chen [39] have shown that age-associated genes identified using bulk expression profiles of the lung are enriched among those induced or suppressed by SARS-CoV-2 infection. However, they did not perform analysis with systems-level approach. Our analysis highlighted RAS and JAK/STAT pathways to be enriched for genes with stronger influence in old AT2 cells and genes up-regulated in COVID infected lung. Ras/MAPK signalling is considered essential for self-renewal of AT2 cell [33]. Similarly, JAK/STAT pathway is known to be activated in the lung during injury [40] and influence surfactant quality[32]. We have used murine aging-lung scRNA-seq profiles however our analysis provides an important insight that regulatory patterns and master-regulators in old AT2 cells are in such a configuration that they could be predisposing it for a higher level of RAS and JAK/STAT signalling. Androgen receptor (AR) which has been implicated in male pattern baldness and increased risk of males towards COVID infection [41] had higher pageRank and degree in old AT2 cells. However, further investigation is needed to associate AR with severity on COVID infection due to ageing. On the other hand, in young AT2 cells, we find a high influence of genes involved in Histamine H1 receptor-mediated signalling, which is known to regulate allergic reactions in lungs [42]. Another benefit of our approach of analysis is that it can highlight a few specific targets of further study for therapeutics. Such as a kinase that binds and phosphorylates c-Jun called as JNK is being tested in clinical trials for pulmonary fibrosis [43]. Androgen deprivation therapy has shown to provide partial protection against SARS-COV-2 infection [44]. On the same trend, our analysis hints that Etv5 could also be considered as drug-target to reduce the effect of ageing induced RAS pathway activity in the lung.

## Methods

We used the term noise in gene-expression according to its definition by several researchers such as Raser and O’Shea [12]; as the measured level of variation in gene-expression among cells supposed to be identical. Hence we first made a base-graph (networks) where supposedly identical cells are connected by edges. For every gene we use this base-graph and apply graph-wavelet transform to get an estimate of variation of its expression in every sample (cells) with respect to other connected samples at different levels of graph-spectral resolution. For this purpose, we first calculated distances among samples (cells). To get a better estimate of distances among samples (cells) one can perform dimension reduction of the expression matrix using tSNE [45] or principal component analysis. We considered every sample (cell) as a node in the graph and connected two nodes with an edge only when one of them was among K-nearest neighbors of the other. Here we decide the value of K in the range of 10-50, based on the number of samples(cells) in the expression data-sets. Thus we calculated the preliminary adjacency matrix using K-nearest neighbours (KNN) based on euclidean distance metric between samples of the expression matrix. We used this adjacency matrix to build a base-graph. Thus each vertex in the base-graph corresponds to each sample and edge weights to the euclidean distance between them.

The weighted graph *G* built using KNN based adjacency matrix comprises of a finite set of vertices *V* which corresponds to cells (samples), a set of edges *E* denoting connection between samples (if exist) and a weight function which gives non-negative weighted connections between cells (samples). This weighted matrix can also be defined as a *NXN* (*N* being number of cells) weighted adjacency matrix *A* where *A_ij_* is 0 if there is no edge between cells *i* and *j*, otherwise *A_ij_* = *weight*(*i, j*) if there exist an edge between *i, j*. The degree of a cell in the graph is the sum of weights of edges incident on that cell. Also, diagonal degree matrix *D* of this graph comprises of degree *d*(*i*) if *i* = *j*, 0 otherwise. A non-normalized graph Laplacian operator *L* for a graph is defined as *L* = *D* - *A*. The normalized form of graph Laplacian operator is defined as:

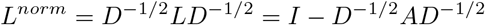

Both laplacian operators produce different eigen-vectors [46]. However, we have used a normalized form of Laplacian operator for the graph between cells. The graph Laplacian is further used for Graph Fourier transformation of signals on nodes (see supplementary Methods) ([47] [46]).

For filtering in the Fourier domain, we used Chebyshev-filter for gene expression profile. We took the expression of each gene at a time considering it as a signal and projected it onto the raw graph (where each vertex corresponds to each sample) object [17]. We took forward Fourier transform of signal and filtered the signal using Chebyshev filter in the Fourier domain and then inverse transformed the signal to calculate filtered expression. This same procedure was repeated for every gene. This would finally give us filtered gene expression.

### Spectral Graph Wavelet Transform

Spectral graph wavelet entails choosing a non-negative real-valued kernel function which can behave as a bandpass filter and is similar to Fourier transform. The re-scaled kernel function of graph laplacian gives wavelet operator which eventually produce graph wavelet coefficients at each scale. However, using continuous functional calculus one can define a function of self adjoint operator on the basis of spectral representation of graph. Although for a graph with finite dimensional Laplacian, this can be achieved by eigenvalues and eigenvectors of laplacian *L* [47]. The wavelet operator is given by *T_g_* = *g*(*L*). *T_g_f* gives wavelet coefficients for a signal *f* at scale = 1. This operator operates on eigenvectors *U_l_* as *T_g_U_l_* = *g*(*λ_l_*)*U_l_*. Hence, for any graph signal, operator *T_g_* operates on the signal by adjusting each graph Fourier coefficient as

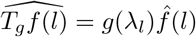

and inverse Fourier transform given as

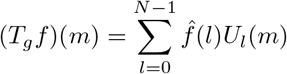

The wavelet operator at every scale *s* is given as 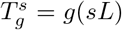. These wavelet operators are localized to obtain individual wavelets by applying them to *δ_n_*, with *δ_n_* being a signal with 1 on vertex *n* and zero otherwise [47]. Thus considering coefficients at every scale, the inverse transform can be obtained as

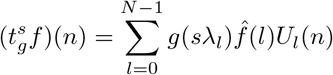

Here, in spite of filtering in Fourier domain, we took wavelet coefficients of each gene expression signal at different scales. Thresholding was applied on each scale to filter wavelet coefficients. We applied both hard and soft thresholding on wavelet coefficients. For soft thresholding, we implemented well-known methods Sure Shrink and Bayes Shrink.

### Choosing threshold for graph-wavelet coefficients

Finding an optimal threshold for wavelet coefficients for denoising linear-signals and images has remained a subject of intensive research. We evaluated both soft and hard thresholding approaches and tested an information-theoretic criterion known as the minimum description length principle (MDL). Using our tool GWNet, user can choose from multiple options of finding threshold such as visuShrink, sureShrink and MDL. Here, we have used hard-thresholding for most the data-sets as proper soft-thresholding of Graph-wavelet coefficient is itself a topic of intensive research and may need further fine-tuning. One can also use hard-threshold value based on the best overlap among predicted gene-network and protein-protein interaction (PPI). While applying it on multiple data-sets we realized that threshold cutoffs estimated by MDL criteria and best overlap of predicted network with known interaction and PPI, were in the range of 60-70 percentile. For comparing predicted network from multiple data-sets, we needed uniform percentile cutoff to threshold graph-wavelet coefficients. Hence for uniform analysis of several data-sets, we have set the default threshold value of 70 percentile. Hence in default mode, wavelet coefficient with absolute value less than 70 percentile was made equal to zero.

### Methods used to infer network among genes

GWNet tool is flexible, and any network inferences method can be plugged in it for making regulatory inferences using a graph-theoretic approach. Here, for single-cell RNA-seq data, we have used gene-expression values in the form of FPKM (fragments per kilobase of exon model per million reads mapped). We pre-processed single-cell gene expression by quantile normalization and log transformation. To start with, we used spearman and Pearson correlation to achieve a simple estimate of the measure of inter-dependencies among genes. We also used ARACNE (Algorithm for the Reconstruction of Accurate Cellular Networks) to infer network among genes. ARACNE first computes mutual information for each gene-pair. Then it considers all possible triplet of genes and applies the Data Processing Inequality (DPI) to remove indirect interactions. According to DPI, if gene i and gene j do not interact directly with each other but show dependency via gene k, the following inequality hold

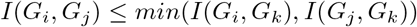

Where *I*(*G_i_, G_j_*) represents mutual information between gene *i* and gene *j*. ARACNE also removes interaction with mutual information less than a particular threshold eps. We have used eps value to Recently Skinnider et al., [6] showed superiority of two measures of proportionality *rho*(*ρ*) and *phi*(*ϕ_s_*) [48] for estimating gene-coexpression network using single-cell transcriptome profile. Hence we also evaluated the benefit of graph-wavelet based denoising of gene-expression with measures of proportionality *ρ* and *ϕ_s_*. The measures of proportionality *ϕ* can be defined as

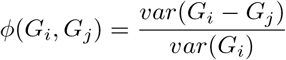

Where *G_i_* is the vector containing log values of expression of a gene *i* across multiple samples (cells) and var() represents variance function. The symmetric version of *ϕ* can be written as

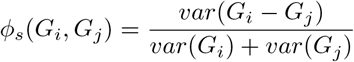

Whereas rho can be defined as

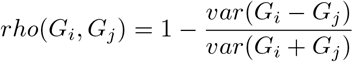

To estimate both measures of proportionality, *ρ* and *ϕ*, we used ‘propr’ package2.0 [49].

### Comparison of Raw and Filtered Graph

The networks inferred from filtered and unfiltered gene-expression were compared to the ground truth. Ground truth for DREAM5 challenge data-set was already available while for single-cell expression, we assembled the ground truth from HIPPIE (Human Integrated Protein-Protein Interaction Reference) Database [50]. We considered all edges possible in network, sorted them based on the significance of edge weights. We calculated the area under the Receiver operator curve for both raw and filtered networks by comparing against edges in the ground truth. Receiver operator is a standard performance evaluation metrics from the field of machine learning, which has been used in the DREAM5 evaluation method with some modifications. The modification for Receiver operating curve here is that for X-axis instead of false-positive rate, we used a number of edges sorted according to their weights. For evaluation all possible edges sorted based on their weights in network are taken from the gene-network inferred from filtered and raw graphs. We calculated improvement by measuring fold change between raw and filtered scores.

### Comparison with other methods

We compared the results of our approach of graph-wavelet based denoising with other methods meant for imputation or reducing noise in scRNA-seq profiles. For comparison we used Graph-Fourier based filtering [17], MAGIC[20], scImpute [21], DCA [22], SAVER [23], Randomly [24], KNN-impute [25]. Brief descriptions and corresponding parameters used for other methods are written in supplementary Method.

### Data Sources

The bulk gene-expression data used here evaluation was download from DREAM5 portal (http://dreamchallenges.org/project/dream-5-network-inference-challenge/). The single-cell expression profile of mESC generated using different protocols [18] was downloaded for GEO database (GEO id: GSE75790). Single-cell expression profile of pancreatic cells from individuals with different age groups was downloaded from GEO database (GEO id:GSE81547). The scRNA-seq profile of murine aging lung published by Kimmel et al. [14] is available with GEO id: GSE132901. While aging lung scRNA-seq data published by Angelids et al. [51] is available with GEO id: GSE132901.

## Availability

The code for graph-wavelet based filtering of gene-expression is available at http://reggen.iiitd.edu.in:1207/GraphWavelet/index.html. The codes are present at https://github.com/reggenlab/GWNet/ and supporting files are present at https://github.com/reggenlab/GWNet/tree/master/supporting$_$files.

## ACKNOWLEDGEMENTS

We thank Dr Gaurav Ahuja for providing us valuable advice on analysis of single-cell expression profile of ageing cells.

## Key Points

- We found that graph-wavelet based denoising of gene-expression profiles of bulk samples and single-cells can substantially improve gene-regulatory network inference.
- More consistent prediction of gene-network due to denoising lead to reliable comparison of predicted networks from old and young cells to study the effect of ageing using single-cell transcriptome.
- Our analysis revealed biologically relevant changes in regulation due to aging in lung pneumocyte type II cells, which had similarity with effects of COVID infection in human lung.
- Our analysis highlighted influential pathways and master regulators which could be topic of further study for reducing severity due to ageing.

**Vibhor Kumar** is an assistant professor at IIIT Delhi, India. He is also an adjunct scientist at Genome Institute of Singapore. His interest include Genomics and signal processing.

**Divyanshu Srivastava** completed his thesis on Graph Signal processing for Masters degree at computational biology department in IIIT Delhi, India. He has applied Graph signal processing on protein structures and gene-expression data-sets.

**Shreya Mishra** is a PhD student at computational biology department in IIIT Delhi, India. Her interest include data sciences and Genomics.

